# Multi-omics analysis of *Bacillus. subtilis* spores formed at different environmental temperatures reveal differences at the morphological and molecular level

**DOI:** 10.1101/2023.06.22.546136

**Authors:** Yixuan Huang, Winfried Roseboom, Stanley Brul, Gertjan Kramer

## Abstract

Spore-forming bacteria play an essential role in the food industry and public health. Through sporulation, bacteria can withstand extreme environmental conditions that vegetative cells cannot survive. Although it is well established that the same environmental factors that affect the growth of vegetative cells also profoundly influence sporulation, the mechanisms of how growth conditions affect spore structure and function remain unknown. Prior research has shown that spores prepared at higher temperatures are more heat resistant than those prepared at lower temperatures. The present study examines, both at metabolomic and proteomic levels, the effect of different sporulation temperatures (25, 37 and 42°C) on the small molecule and protein composition of spores (strain PY79) of the model organism *B. subtilis*. Through differential harvesting times, spores of the same developmental stage were obtained for each temperature regime. The heat resistance, dipicolinic acid content, germination kinetics and spore morphology were assayed to compare spore properties. Metabolome and proteome analysis yielded unparalleled broad molecular detail of the spores formed at different environmental temperatures. Our findings indicate that peptidoglycan biosynthesis and 28 outer-layer proteins play a crucial role in the functional diversity of spores produced by *B. subtilis* under varying temperatures.

## Introduction

Spore-forming bacteria, of the species *Bacillus* are of high interest in relation to food spoilage or food poisoning and become common contaminants in food processing and production (Checinska et al., 2015). These stress resistant spores can help vegetative cells to survive from farm to fork throughout the food chain as they facilitate resistance to extreme environments like food-processing operations characterized by ultra-high-pressure, elevated temperatures or ϒ-irradiation. The surviving spores can be dispersed by wind, living hosts or raw materials to anywhere, break dormancy and start a new lifecycle quickly through germination when the environmental conditions are favorable for outgrowth and subsequent vegetative cell growth leading often-times to either infection or toxin production (Bressuire-Isoard et al., 2018). Hence, bacterial spore formers easily cause next to common food spoilage also food borne infection and food poisoning (Gauvry et al., 2021). Prior research has proven that the resistance properties, production, and functions of spores are influenced by sporulation conditions to a great extent. For instance, *Bacillus. subtilis* spores prepared on solid Schaeffer’s-glucose (SG) agar plates showed a significantly thinner outer coat layer and higher heat resistance than those of prepared in liquid MOPS buffered defined medium (Abhyankar et al., 2016). Observations on accumulated mineral concentrations in the spore core suggested that it is influenced to a large extent by the composition of the sporulation medium (Baril et al., 2012). A lower sporulation temperature can result in an increment of the total amount of fatty acid (FA) in *Bacillus. cereus* spores (Planchon et al., 2011). From the above, we learn that the variation in environmental factors not only influences the growth of *Bacillus* cells but also sporulation through intricate developmental processes.

A better and complete understanding of the mechanisms of development and resistance of spores based on environmental factors is essential to find methods that optimally prevent undesired spore survival, germination and outgrowth of vegetative cells. In the food industry, heat treatment is mostly applied to reduce the microbial load on foods, which also provides a positive selection of spore formers due to their above described intrinsic heat resistance (Postollec et al., 2012). However, given the type of food and nutritional considerations, the intensity and pattern of the sterilization process always needs to be balanced against unwanted effects on the organoleptic quality of the food products. Thus, the extent of spore inactivation options by heat only is limited by food quality constraints (Trunet et al., 2015). Furthermore, by controlling the recovery conditions and preventing the germination of surviving spores in processed food one can achieve a better end result at lower thermal input using a so-called hurdle preservation approach (Liu et al., 2022). With respect to recovery, the spores’ properties varied with the sporulation condition and can greatly affect spore germination. Currently, many studies have concentrated on ecological niches of spore-formers (Miller et al., 2015), and the resistance abilities of spores (Mtimet et al., 2015), etc. But few studies pay attention to how metabolic, physiological or molecular adaptations during growth will interfere with sporulation and molecular and physiological mechanisms behind these observations.

In this study, spores of a laboratory strain of the model organism *B. subtilis*, spores produced at three different temperatures (25, 37 and 42°C) were investigated for a variety of physiological parameters as well as their metabolomes and proteomes. 25°C is often encountered in soil and parts of the food chain (Wang et al., 2021), 37°C is an optimal growth and sporulation temperature for Bacillus subtilis generally used in the laboratory setting while 42°C is an environmental stress temperature. First, we confirmed the harvesting time to obtain spores of the same developmental stage grown at the three different temperatures using fluorescent strains to gauge sporulation stage (Isticato et al., 2020) during these different conditions. Subsequently we determined the heat resistance, dipicolinic (DPA) content, speed of initiating of germination and spore morphology to compare physiological properties of these spores. We then used 60% EtOH extraction to enable simultaneous analysis of the metabolome, lipidome and proteome from one single sample. The biosynthesis of peptidoglycan and the involvement of 28 outer-coat layer proteins are crucial factors underlying the functional diversity of spores generated by *B. subtilis* when exposed to different temperatures.

## Materials and Methods

### Strains, sporulation and calibrating spore stage during different sporulation temperatures

The strain *B. subtilis* PY 79 (wild type), RH2466 (pspoIIQ::gfp) (Donadio et al., 2016), RH2467 (pgerE::gfp) (Donadio et al., 2016) and RH238(CotC::gfp) (Isticato et al., 2007) were used in this study. The three isogenic strains of *B. subtilis* carrying green fluorescent protein (gfp) were acquired from Prof. Ezio Ricca’s lab (University of Naples Federico II). Both wild type and mutant strains were cultured and harvested as described previously (Kort et al., 2005). In brief, the selected single colony from a LB agar plate was inoculated in LB medium and then transferred the log phase bacteria to MOPS medium, with 40 mM glucose and 40 mM NH_4_Cl, to make a series of dilutions cultured at 25, 37 or 42°C overnight. 1 ml dilutions with an OD_600_ of 0.3-0.4 were transferred to 100ml MOPS medium at 25, 37 or 42°C and 200 rpm to induce sporulation. Collected spores were washed three times with sterile cold milliQ-water and purified by Hiztodenz gradient centrifugation (Ghosh et al., 2015). In order to harvest similar age spores, fluorescence microscopy and number of free spores were determined as described previously (Isticato et al., 2020).

### Heat resistance assay

The purified spores from the wild type cultured under 25, 37 or 42°C were subjected to 85°C following a previously established protocol (Abhyankar et al., 2016). 1 ml of purified spores (OD_600_ ∼2) were incubated in a water bath at 70°C for 30 min to kill vegetative cells. Spores were then placed on ice for 15 min and then injected with a syringe into metal screw-cap tubes containing 9 ml of sterile Milli-Q water which was heated for 30 min in a glycerol bath (85°C). After that, the metal tubes were cooled on ice for 15 min. From this heat challenged spore suspensions, a series of dilutions were made and spread on LB agar plates for counting the number of colonies. Three biological replicates were performed.

### DPA Measurement

The DPA measurement protocol was based on a modification of a previously published method (Donnelly et al., 2016). 1 ml of spores with an OD_600_ of 2 from each strain was suspended in buffer 1 containing 0.3 mM (NH_4_)_2_SO_4_, 6.6 mM KH_2_PO_4_, 15 mM NaCl, 59.5 mM NaHCO_3_, and 35.2 mM Na_2_HPO_4_. The suspended spores were heated at 100° C for 1h to induce a leakage of DPA with spores held at 37° C as a negative control. Samples were then centrifuged at 15,000 rpm for 10 minutes. 10 µL of the resulting supernatant was mixed with 115 µL of buffer 2 (1 mM Tris, 150 mM NaCl) both with and without 0.8 mM terbium chloride (Sigma-Aldrich, St. Louis, MI, USA). Following a 15-minute incubation, the fluorescence was measured on Synergy Mx microplate reader (BioTek, Bad Friedrichshall, Germany) with an excitation wavelength of 270 nm and a reading wavelength of 545 nm. The background fluorescence was subtracted from all the samples, and a calibration curve of 0–125 µg/mL DPA (Sigma-Aldrich, St. Louis, MI, USA) was used to calculate DPA concentrations. Three biological replicates were used for all conditions.

### Live imaging of spores

The purified spores were immobilized on 1.5% agarose pads with complete minimal medium with a 1.5 cm × 1.6 cm Gene Frame® (Thermo Scientific, Landsmeer, Netherlands) attached to a microscope slide. The germination was induced by adding 10 mM L-valine and AGFK (10 mM L-asparagine, 10 mM glucose, 1 mM fructose and 1 mM potassium chloride), and the time-lapse phase contrast images were captured by a Nikon Eclipse Ti, which was coupled a Prior Brightfield LED, a Nikon CFI Plan Apo Lambda 100X Oil, C11440-22CU Hamamatsu ORCA flash 4.0 camera, LAMBDA 10-B Smart Shutter from Sutter Instrument, an OkoLab stage incubator, and NIS elements software version 4.50.00. A sample frequency of 1 frame per 1 min was recorded for the individual spores for 6 hours. Spores of each sporulation temperature from three biological replicates were imaged at 37°C to track spore germination in detail. The obtained living images were analyzed with the modified ImageJ macro SporeTrackerX (Omardien et al., 2018), which can assess germination events based on the changes of spore brightness (Pandey et al., 2013).

### Morphological observation

The similar developmental stage spores induced under 25, 37 or 42°C were harvested, to observe their morphology and calculate their aspect ratio as described before (Mou et al., 2020). Phase contrast images were captured by collecting purified spores (100 μL), resuspending the spores in 10μl of 1× PBS buffer and observing 1 μL with a Nikon cell observer using 100× Oil objective Nikon CFI Plan Apo Lambda. For each imaging process, 7-10 fields of views were captured. To analyze the aspect ratio of spores, ImageJ/Fiji (http://fiji.sc/Fiji) was applied for measurement. For this analysis, 1000 spores for every different temperature (25, 37 or 42° C) sample were used.

### Metabolite extraction

For metabolite extraction of spore samples, 20OD spore sample for each replicate was homogenized in 3mL ice-cold 60% ethanol aqueous solution with Zirconium-silica beads (0.1 mm, BioSpec Products, Bartlesville, OK, USA) in a screw-top tube. After that, transferring the supernatant a new tube and washing the beads three times using ice-cold 60% ethanol. The washing solutions were combined with the previous supernatant and centrifuged for 15 min at 4°C, 8000 rpm. The obtained supernatant and pellet were dried under a speed vacuum concentrator. The dried metabolites and pellets were kept at −80 °C until LC-MS analysis. Quality control (QC) samples were prepared by pooling 10 µL homogenate of each spore sample.

### Single Tube Solid Phase Sample Preparation (SP3)

20 μg/μl beads (magnetic carboxylate modified) were prepared at room temperature and the protein pellet from the 60% EtOH extraction dissolved in 300 μL 1% SDS (made by 100mM Ambic). Prior to reduction of disulfide bridges by 10mM Tris(2-carboxyethyl) phosphine hydrochloride (TCEP) and carbamidomethylation of free cysteines by 30mM 2-Chloroacetamide (CAA), a bicinchoninic acid (BCA) assay was performed according to the manufacturer’s protocol using the Pierce™ BCA Protein Assay Kit (CAT NO. 23250) to determine the protein concentration. Subsequently 20 μg of protein was mixed with 2 μL prepared bead solution in a PCR tube and samples were processed as described previously (Hughes et al., 2014). Trypsin (protease/protein, 1:50, w/w) was added for protein digestion at 37°C overnight. Following tryptic digestion, formic acid (FA) was added to the supernatant to halt digestion and peptide solutions recovered from the magnetic beads using a magnetic stand for subsequent LC-MS analysis.

### LC-MS/MS analysis

#### Metabolomics analysis

For apolar metabolites analysis, the reconstitution was in 200 μL water and polar metabolites analysis was in 200 μL ACN/ H_2_O (1:1). 10-μl samples were injected for non-targeted metabolomics onto a CSH- C18 column (100 mm × 2.1 mm,1.7 μm particle size, Waters, Massachusetts, USA) for apolar metabolites analysis or a BEH-Amide column (100 mm × 2.1 mm,1.7 μm particle size, Waters, Massachusetts, USA) for polar metabolites by an Ultimate 3000 UHPLC system (Thermo Scientific, Dreieich Germany). Using a binary solvent system (A: 0.1% FA in water, B: 0.1% FA in ACN) metabolites were separated on the CSH-C18 column by applying a linear gradient from 1% to 99% B in 18 minutes at a flow rate of 0.4 ml/min, while metabolites were separated on the BEH-Amide column by applying a linear gradient from 99% B to 40% B in 6 minutes and then to 4% B in 2 minutes at a flow rate of 0.4 ml/min. Eluting analytes were electrosprayed into a hybrid trapped-ion-mobility-spectroscopy quadrupole time of flight mass spectrometer (tims TOF Pro, Bruker, Bremen Germany), using a capillary voltage of 4500 volt in positive mode and 3500 volt in negative mode, with source settings as follows: end plate offset 500 volt, dry temp 250 °C, dry gas 8 l/min and nebulizer set at 3 bar both using nitrogen gas. Mass spectra were recorded using a data dependent acquisition approach in the range from m/z 20-1300 for polar and 100-1350 for the apolar metabolites in positive and negative ion mode using nitrogen as collision gas. Auto MS/MS settings were as follows: Quadrupole Ion Energy 5 eV, Quadrupole Low mass 60 m/z, Collision Energy 7 eV. Active exclusion was enabled for 0.2 min, reconsidering precursors if ratio current/previous intensity > 2.

#### Proteomics analysis

Peptides were dissolved in 6 μL water containing 0.1% FA and 3% ACN and then 200ng/μl (measured by a NanoDrop at a wavelength of 215 nm) of the peptide was injected by an Ultimate 3000 RSLCnano UHPLC system (Thermo Scientific, Germeringen, Germany). Following injection, the peptides were loaded onto a 75um x 250 mm analytical column (C18, 1.6 μm particle size, Aurora, Ionopticks, Australia) kept at 50°C and flow rate of 400 nl/min at 3% solvent B for 1 min (solvent A: 0.1% FA, solvent B: 0.1% FA in ACN). Subsequently, a stepwise gradient of 2% solvent B at 5 min, followed by 17% solvent B at 24 min, 25% solvent B at 29 min, 34% solvent B at 42 min, 99% solvent B at 33 min held until 40 min returning to initial conditions at 40.1 min equilibrating until 58 min. Eluting peptides were sprayed by the emitter coupled to the column into a captive spray source (Bruker, Bremen Germany) which was coupled to a TIMS-TOF Pro mass spectrometer. The TIMS-TOF was operated in PASEF mode of acquisition for standard proteomics. In PASEF mode, the quad isolation width was 2 Th at 700 m/z and 3 Th at 800 m/z, and the values for collision energy were set from 20-59 eV over the TIMS scan range. Precursor ions in an m/z range between 100 and 1700 with a TIMS range of 0.6 and 1.6 Vs/cm^2^ were selected for fragmentation. 10 PASEF MS/MS scans were triggered with a total cycle time of 1.16 seconds, with target intensity 2e^4^ and intensity threshold of 2.5e^3^ and a charge state range of 0-5. Active exclusion was enabled for 0.4 min, reconsidering precursors if ratio current/previous intensity >4.

### Data processing and bioinformatics

The metabolites mass spectrometry raw files were submitted to MetaboScape 5.0 (Bruker Daltonics, Germany) used to perform data deconvolution, peak-picking, and alignment of m/z features using the TReX 3D peak extraction and alignment algorithm (EIC correlation set at 0.8). All spectra were recalibrated on an internal lockmass segment (NaFormate clusters) and peaks were extracted with a minimum peak length of 8 spectra (7 for recursive extraction) and an intensity threshold of 1000 counts for peak detection. In negative mode ion deconvolution setting, [M-H]- was set for primary ion, seed ion was [M+Cl]- and common ions were [M-H-H_2_O]-, [M+COOH]-. For positive mode, the primary ion was [M+H] +, seed ions were [M+Na] +, [M+K] +, [M+NH_4_] + and [M-H-H_2_O] + was the common ion (Edwards-Hicks et al., 2020). Features were annotated, using SMARTFORMULA (narrow threshold, 3.0 mDa, mSigma:15; wide threshold 5.0 mDa, mSigma:30), to calculate a molecular formula. Spectral libraries including Bruker MetaboBASE 3.0, Bruker HDBM 2.0, MetaboBASE 2.0 in silico, MSDIAL LipidDBs, MoNA VF NPL QTOF, AND GNPS export, were used for feature annotation (narrow threshold, 2.0 mDa, mSigma 10, msms score 900, wide threshold 5.0 mDa, mSigma:20 msms score 800). Analyte listS containing 667 compounds with a retention time (RT) (narrow threshold, 1.0 mDa, 0.05 minutes, mSigma: 10, msms score 900; wide threshold 5.0 mDa, 0.1 min, mSigma 50, msms score 700) was also used to annotate deconvoluted features. An annotated feature was considered to be of high confidence if more than two green boxes were present in the Annotation Quality column of the program and low confidence if less than two green boxes were present. Resulting data were exported for further analysis with MetaboAnalyst 5.0 (https://www.metaboanalyst.ca/MetaboAnalyst/home.xhtml). At first, the data showing a poor variation were filtered on Inter Quartile Range (IQR) and then features were normalized by median normalization, scaled by Auto scaling and transformed to a logarithmic scale (base of 2).

Generated proteomics raw data was searched in Maxquant (Cox and Mann, 2008) (version: 1.6.14.0) and were matched to an UniProt proteome database (proteome ID UP000001570). Methionine oxidation was set as variable modifications, whereas cysteine carbamidomethyl was set as a fixed modification with a maximum of 2 missed cleavages of Trypsin/P. The “Match between runs” was selected with a matching time window of 0.2 minutes and TIMS-DDA was chosen for type of group. Label-free quantification (LFQ) was performed using classic normalization and a minimum ratio count of 2. All the other setting parameter was default. The quantified protein data file was reported in the output proteinGroup.txt, which were further analyzed and visualized using Perseus (version 1.6.15.0) (Tyanova et al., 2016). Data were filtered to remove potential contaminants, reverse hits, protein groups “only identified by site” and the number of unique peptides more than 1 were kept.

The differentially expressed proteins and metabolites were determined by using R/ Bioconductor package *limma* (Ritchie et al., 2015). R package Mfuzz (Kumar and E. Futschik, 2007) was applied for cluster and visualization.

## Results

### Determination of time to harvest spores of similar developmental stage at 25, 37 or 42°C

To initially assess the degree of delay in sporulation of spores cultured at different temperatures, 500 μL aliquots were collected every 1.5 h to check with a wide-field fluorescence microscope to record the time of appearance of the GFP fluorescent signals from reporter proteins (see Materials and Methods) at the three growth temperatures. Table 1 shows that the green fluorescence appeared at different time points, especially for the sporangia produced at 25°C, while a slight acceleration was observed at 42°C compared with cells growing at 37°C. The time of appearance of GFP fluorescentce at 25°C was divided by that at 42°C and the resulting ratio was considered as the sporulation delay factor. For the GFP fluorescent protein controlled by the σF, σK or σK-GerE dependent promoters, 1,62, 1.72 and 1.68 were calculated respectively and the mean value was 1.67 (Supplementary Table S1). Because there were only minimal differences on the delay factor observing between cells at 37 versus 42 °C, it was not considered as different in our further study. The percentage of free spores producing at 42, 37 or 25℃ was determined at different time points to monitor the sporulation under the phase-contrast microscope. We selected 85% of free spores as a standard to further determine the time to harvest similar developmental stage spores. Finally, 48 h at 37 and 42°C and 91 h at 25°C were chosen as time of harvest and shown in supplementary Fig. S1. These two experiments were in good agreement with each other and 48 h at 37 and 42°C and 91 h at 25°C were used for all subsequent experiments.

### Spore properties and morphology analysis

To assess the wet-heat resistance of spores produced at 25, 37 and 42°C, similarly staged *B. subtilis* PY 79 spores were heat treated at 85°C. Clearly, 85°C treatment led to statistically significant decrease (p < 0.05) in surviving spores of those produced at 25°C compared with those produced at 37 or 42°C (Fig. 1A). However, there were no significant differences observed in surviving spores between 37 or 42°C. DPA levels were significantly higher in *B. subtilis* spores produced from cells sporulating at 42°C compared to those produced upon sporulation at 25°C (Fig. 1B). Furthermore, the time required to start germination by using AGFK and valine as germinants was observed for individual spores. Around 125 spores for each temperature were analyzed with SporeTrackerX. *B. subtilis* PY79 spores produced at temperatures of 37°C and 42°C exhibit a significant difference in their germination onset time when compared to those produced at 25° C. Fig. 1C showed that 25° C spores need less time to start germination compared to 37 and 42° C spores, with no significant differences between 37 and 42° C. For morphology analysis, 1000 individual spores of each sample were measured on aspect ratio. Phase contrast images of spores at various temperatures are presented in Fig. 1D, revealing distinguishable differences in spore morphology. Fig. 1E provided the corresponding statistical analysis on the aspect ratio of spores, which indicates an increasing trend in aspect ratio as the sporulation temperature rises. The spores undergoing sporulation at 25°C show a predominantly spherical morphology, while those generated at 37°C and 42°C exhibit an elongated rod-like shape. Generally, spores produced at 37°C and 42°C contain a higher concentration of DPA, which might contribute to their enhanced resistance to heat treatment. In contrast, spores produced at 25°C demonstrate a slightly faster germinant response when compared to spores produced at 37°C and 42°C and are significantly more thermolabile.

**Fig. 1.**
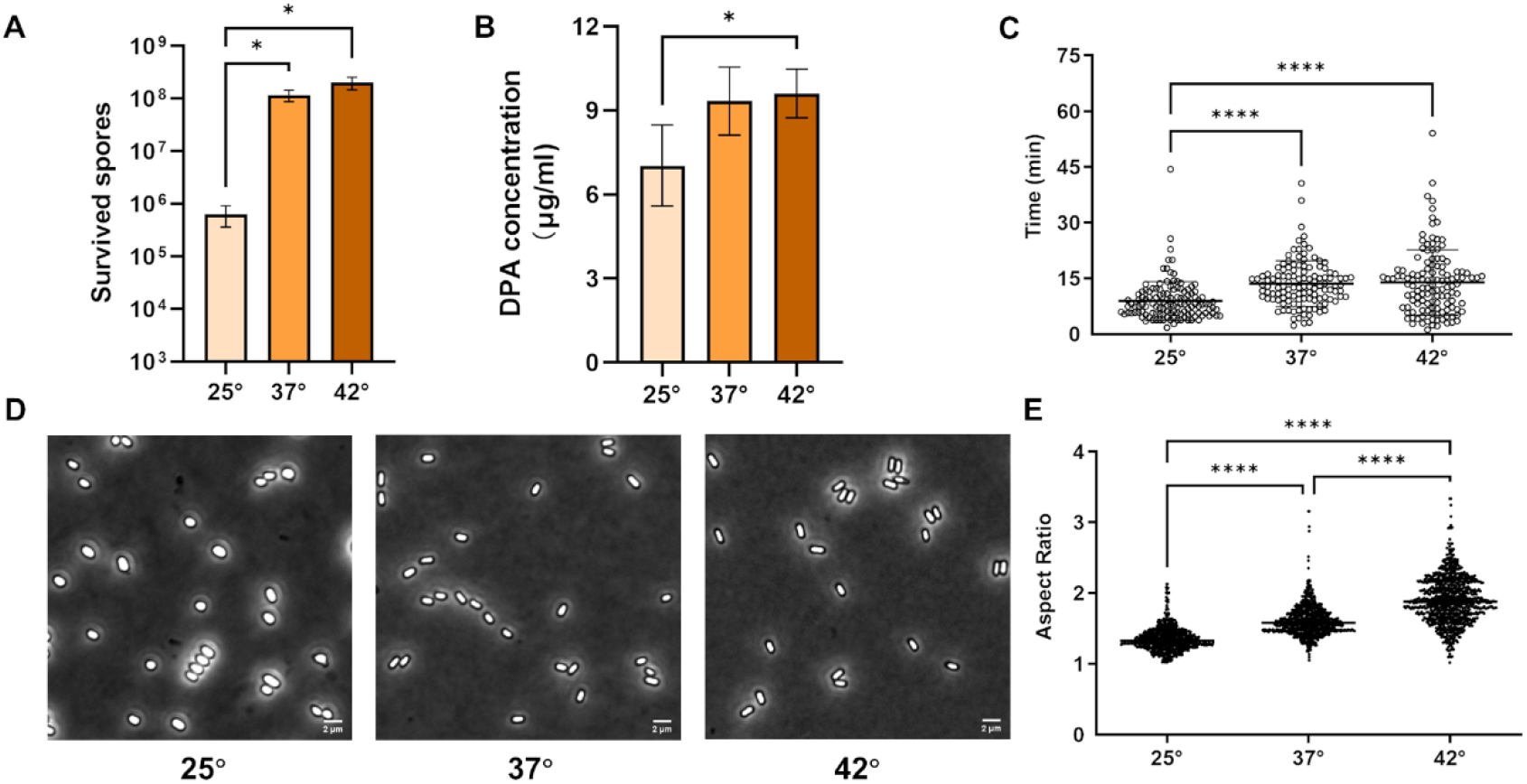
Effects of sporulation temperatures on spore wet heat resistance (A), DPA content (B), germination efficiency (C), morphology (D) and corresponding statistical analysis of shape through aspect ratio (E). * p < 0.05, ** p < 0.01, *** p < 0.001, **** P < 0.0001.

### Proteomic analysis

#### Global spore proteome changes with sporulation temperature

High-throughput quantitative proteomics was utilized to investigate the global proteome changes in spores formed during 25, 37, or 42 °C exposure. In total, 1281 proteins were quantified with over 70% data completeness across all the groups (Supplementary Table S2). We sought to investigate how sporulation temperature affected the protein composition of the formed spores. The Limma package was employed to determine the differential proteins between spores produced at 42°C and those produced at 25°C and 37°C with the criteria of an adjusted p-value of less than 0.05 and a fold change greater than 1.5. It was found that 442 proteins were significantly altered in abundance at 42°C compared to 25°C, and 253 proteins at 42°C compared to 37°C (Fig. 2A and B, Supplementary Table S3). For the comparison of 25°C vs. 37°C, the data can be found in Fig. S2 A and B. To further investigate the impact of sporulation temperature on protein composition of the resulting spores, a Venn diagram was generated using the differentially expressed proteins from each temperature comparison. The analysis revealed a total of 157 proteins (29%) that were shared between the 42°C vs. 25°C and 42°C vs. 37°C comparisons (Fig. 2C). . Within the subset of shared proteins, 64 were found to be up-regulated and 90 were down- regulated in response to the higher temperature compared to the lower temperature, with only three proteins exhibiting opposite patterns of regulation, as indicated in the table in Fig. 2D. Out of the 157 shared proteins, a subset of 154 proteins with the same pattern of regulation were annotated with their functional categories from SubtiWiki. This functional annotation showed the top three relevant categories with the largest changes to be sporulation, protein synthesis, modification and degradation, and membrane proteins (Fig. 2E).

**Fig. 2.**
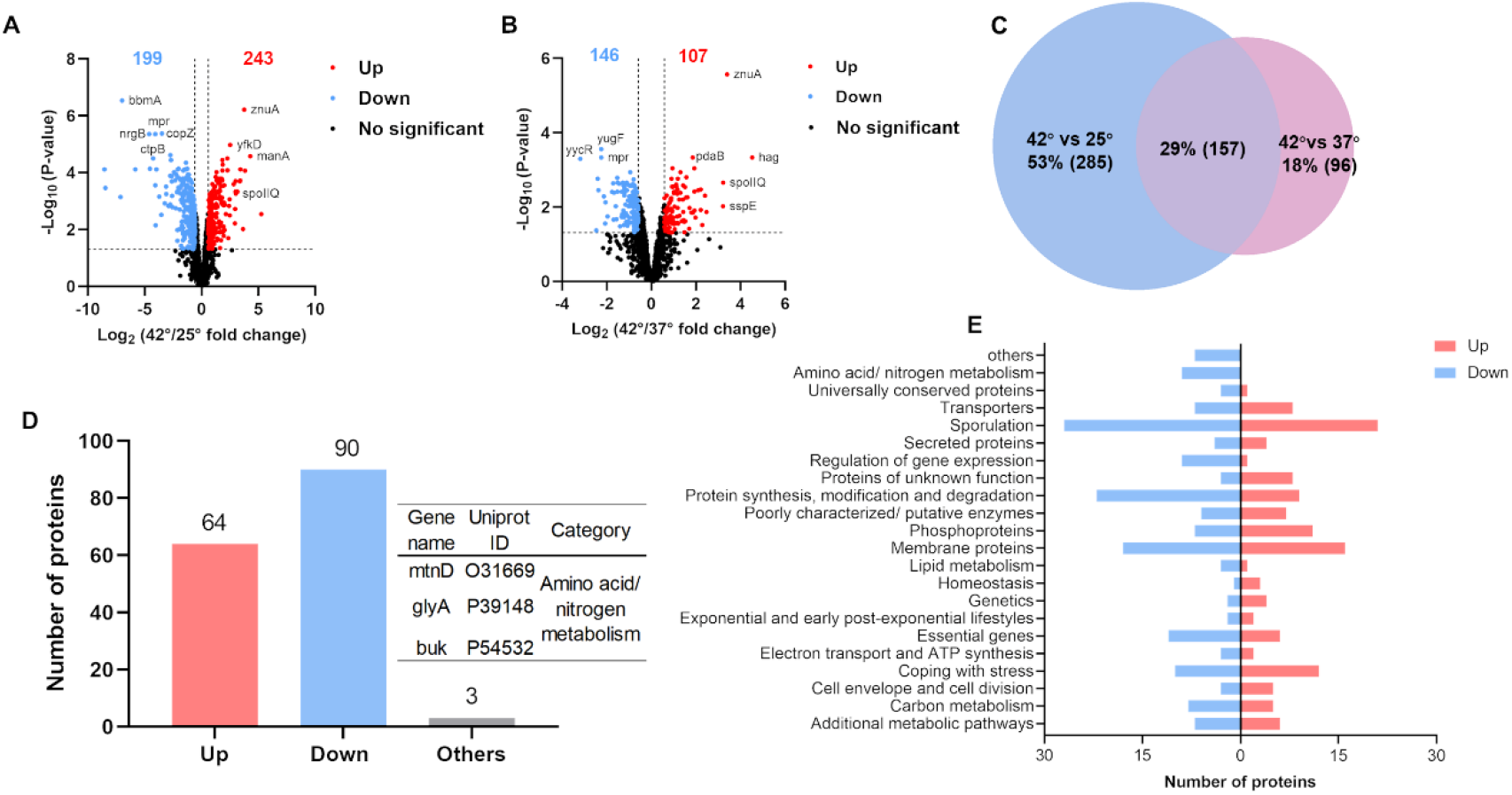
Analysis of proteomics profiles of *B. subtilis* spores sporulated at 42, 37 and 25℃. The volcano plots show 42 vs. 25℃ (A) and 42 vs. 37℃ (B). Venn diagram of differentially expressed proteins based on the comparisons in the volcano plots (C). No of common proteins that show altered temperature dependent levels in dormant spores formed at different temperatures (D). Categorical classification of common differentially expressed proteins according to SubtiWiKi (E).

#### Sporulation temperature induces dynamics changes in proteome and regulators

To further investigate the temperature regulation and functional changes during sporulation, a soft-clustering analysis was conducted using Mfuzz to analyze the differentially expressed proteomes. The analysis focused on clusters exhibiting changes in response to temperature variations during sporulation. Fig. 3A displays a 4-clustered heatmap of log-transformed data that were assembled using Z-score normalization and Fig. 3B further illustrates each distinct patterns of protein expression in response to temperature changes during sporulation. Cluster 1 and 2, comprising 147 and 104 proteins, respectively, exhibited an increase in abundance of proteins in response to higher temperatures. Distinct patterns of protein expression were observed in cluster 2 and cluster 3, which suggested that there may be a coordinated response of proteins involved in sporulation to changes in temperature. KEGG pathway enrichment analysis for each cluster is shown in Supplementary Fig. S3. Sporulation proceeds in a sequential manner under the regulation of the specific sigma factors σF, σE, σG, and σK, which activate compartment-specific transcriptional programs that guide the morphological stages of sporulation (Fimlaid and Shen, 2015; Hilbert and Piggot, 2004). To investigate the impact of temperature on regulators of sporulation stage, we assessed the protein abundance in the resulting spores for proteins under the control of these sigma factors (Fig. 3C). The protein expression level under the regulation of σF significantly increased, whereas those under σE and σG showed only slight increases with increasing temperature. In the case of the σK module, most of the relevant protein levels exhibited a dramatic decrease in spores formed at 42°C.

**Fig. 3.**
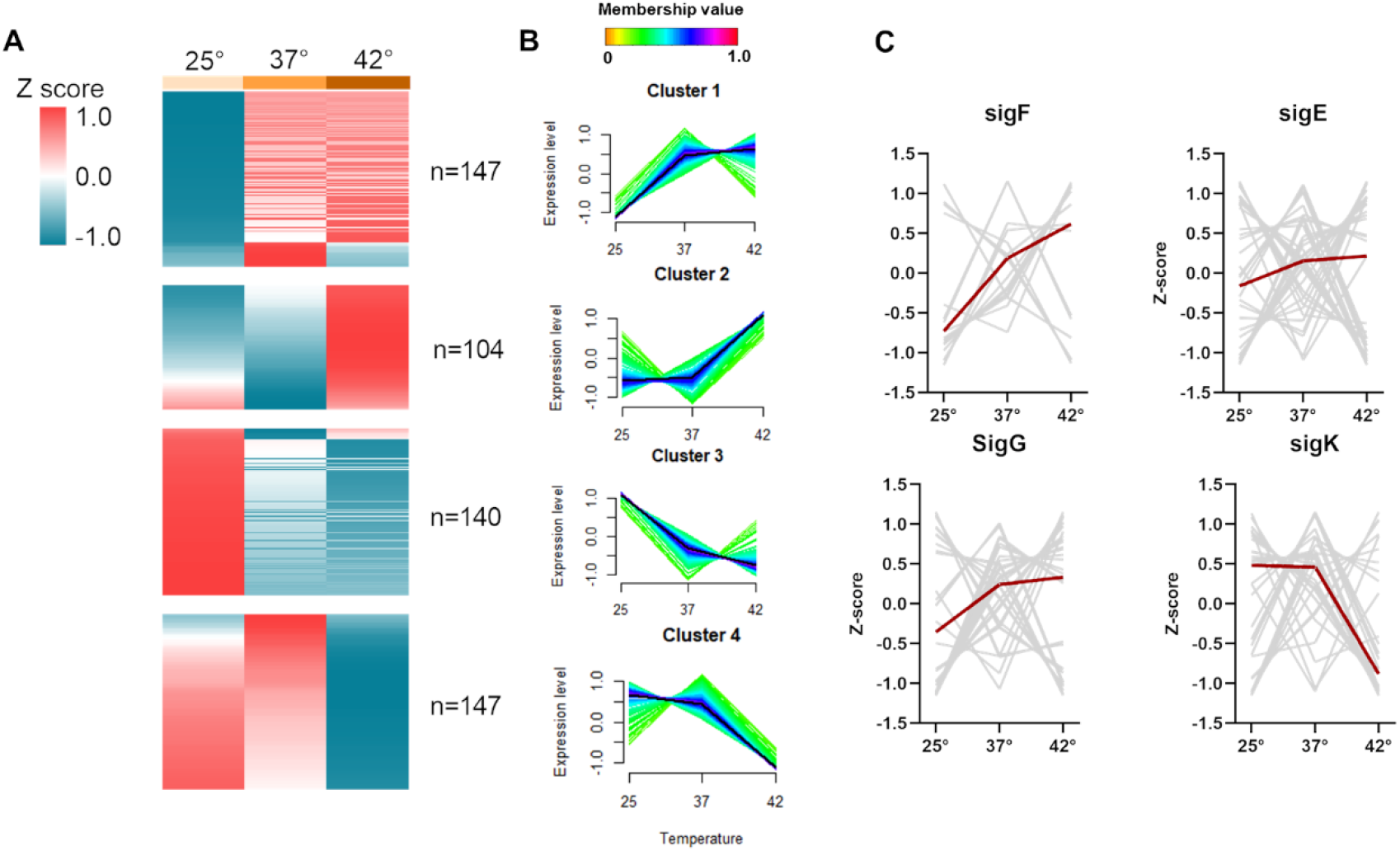
Dynamics of protein composition induced by varying sporulation temperature. Heat map shows the expression level of proteins under each temperature (A), Mfuzz c-means clustering grouped expression profiles of proteins in relation to sporulation temperature (B), examination of differential temperature dependent accumulation of proteins under control of different sigma factors during sporulation (C).

#### Comparison of spore specific proteins

The acquisition of spore resistance to extreme conditions is influenced by several factors, including unique coat and inner core structures, small proteins that saturate DNA (Setlow, 2014). Accordingly, we analysed small acid-soluble spore proteins (SASPs), cell shape proteins, coat proteins and germination proteins to investigate their levels in spores related to sporulation temperature. Our findings revealed a notable increase in the abundance of three SASPs, sspJ, sspE, and sspA, in spores produced at 42°C (Fig. 4A). This observation may be associated with the spores’ enhanced heat resistance, along with the higher concentration of DPA in the spore core. In addition, four heat shock proteins (HSPs) DnaK, GroES, HtpG, and GroEL, exhibited significant higher levels at 42°C (Fig. 4B). However, DisA displayed a inverse trend, with lower levels in spores formed at the higher temperature. In addition, RodZ, YubA, RpeA, Mbl and SsdC which are involved in determining cell and spore shape are less abundant in 25°C spores (Fig. 4C) which could be linked to the changed morphology observed for these spores (Fig 1d) . As depicted in Fig. 4D, a heatmap representing 28 coat proteins grouped into two unsupervised clusters. Notably, cluster 1, containing 18 proteins, such as SpoVM, CotQ, CotB, and CotSA, exhibited considerably lower abundance in spores produced at 42°C, while cluster 2, comprising 10 proteins, showed significantly higher abundance compared to spores produced under 25°C and 37°C. Spores produced upon sporulation at 42°C had decreased protein levels of GerE and Ykvu, but exhibited elevated levels of seven other germination-related proteins, such as GerQ (Fig. 4E).

**Fig. 4.**
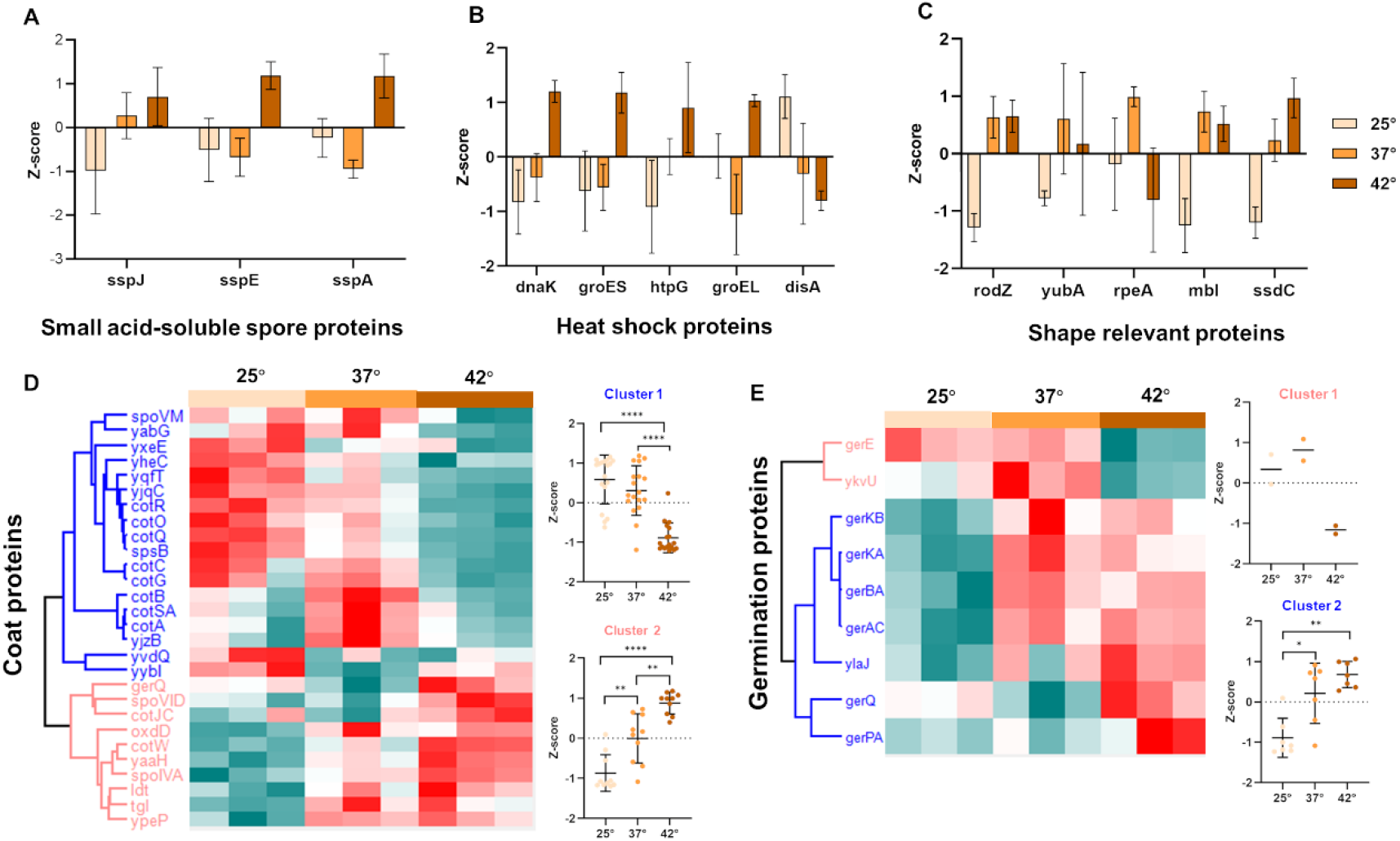
Variation of spore function protein abundance in spores formed at 42, 37 or 25℃. Bar plots and heatmaps showing the protein levels of SASPs (A), HSPs (B), shape relevant proteins (C), coat proteins (D) and germination proteins (E) in spores formed at different environmental temperatures. * p < 0.05, ** p < 0.01, *** p < 0.001, **** P < 0.0001.

### Metabolomics analysis

#### Changes of the spore metabolome composition induced by sporulation temperature

To understand how sporulation temperature affects the spore metabolome, similar stage spores formed upon sporulation at different temperatures were harvested for untargeted metabolomics analysis. The quality assessment of the analysis was conducted through quality control (QC) samples and is described below. The coefficient of variation (CV), with a threshold of less than 0.2, were assessed across all temperature groups and detected metabolites in the QC samples (Supplementary Fig. S4). The percentage of CV< 0.2 were determined to be an average of 78% in three temperature groups and 88% in QC, respectively, indicating good quality for further analysis (Carobene et al., 2013). The current analysis putatively annotated 439 metabolites, with principal-component analysis (PCA) demonstrating a clear separation of the datasets that implied the existence of sporulation temperature-specific metabolic changes (Fig. 5A and Supplementary Table 4). Furthermore, we performed different comparisons of 42℃ versus 37℃ (comparison A) and 42℃ versus 25℃ (comparison B). After filtering with an adjusted p- value <0.05 and fold change >1.5, sporulation temperature was shown to impact on the abundance of 32 and 186 metabolites in Fig. 5B and C (Supplementary Table 5), respectively. For the comparison of 25°C vs. 37°C, the data can be found in Fig. S2 C and D. Overall, more metabolites decreased in comparison A (21) than B (161) upon analyzing the higher temperature against the lower temperature. We used Venn diagrams to identify which metabolites were specifically and most significantly affected by sporulation temperature and aimed at their characterization based on established metabolite classes. Fig. 5D shows 25 (13%) metabolites are shared by these two comparisons while 161(83%) metabolites and 7 metabolites (4%) are unique to the 42℃ versus 25℃ and 42℃ versus 37℃ groups respectively. The production of spores at 42℃ significantly reduced the levels of organic acids and derivatives as well as organic oxygen compounds, while causing a higher presence of PE 15:0-15:0, PG 13:0-15:0, and deoxyadenosine monophosphate (Fig. 5E-H).

**Fig. 5.**
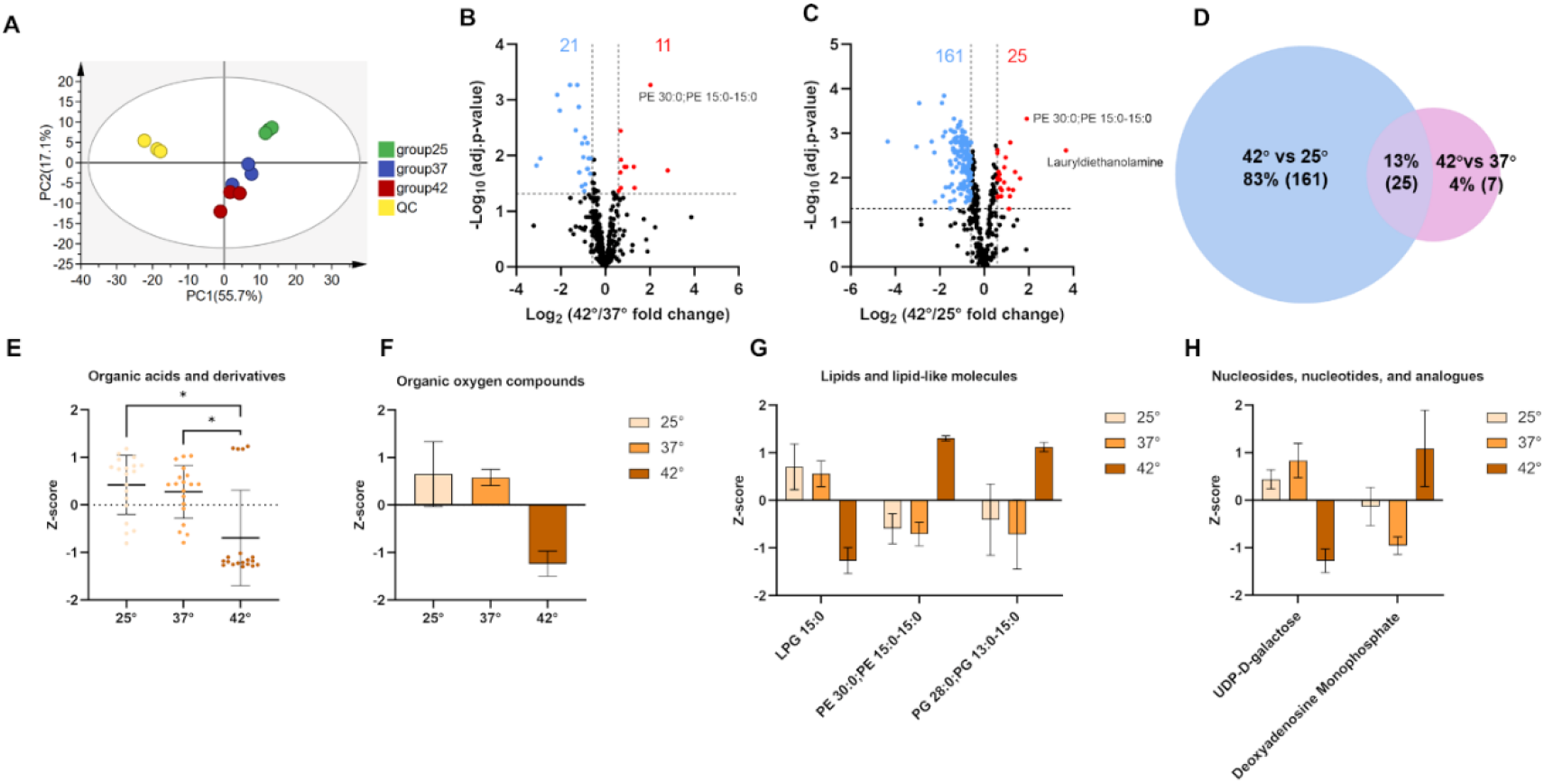
Analysis of metabolomics profiles of *B. subtilis* spores sporulated upon exposure to 42, 37 and 25℃. PCA analysis of all metabolites (A). Volcano plots of metabolites showing to 42 vs. 37℃ (B) and 42 vs. 25℃ (C). Venn diagram of differential metabolite composition based on the comparisons in panels b and c (D). Chemical category classification of common differentially accumulated metabolites in spores formed at different temperatures (E-H).

#### Sporulation temperature induces dynamics changes in the metabolome

We next sought to compare the trend in metabolomic profiles for the different sporulation temperatures. Soft clustering analysis identified 4 metabolite clusters using a total 193 differentially expressed metabolites (Fig. 6A). Analysis of the metabolites in each cluster revealed 4 different profiles according to their membership value (Fig. 6B). Membership values quantify the level of metabolite conformity within its affiliated cluster and high values suggest a pronounced correlation between metabolite levels and the centroid of the cluster (Kumar and E. Futschik, 2007). Different KEGG pathways were enriched in clusters 2–4 (Fig. 6D). Because there was no pathway enriched in cluster 1, the data is not shown. Notably, the metabolite levels in cluster 4 exhibited a linear decrease with increasing temperature, emphasizing a strong relationship between these metabolites and sporulation temperature. Peptidoglycan (PG) biosynthesis was identified as the most significantly enriched pathway in this cluster. The chemical classification composition of different clusters is shown in Fig. 6C. The data indicates that organic acids and derivatives represent a high proportion of the identified mass spectrometric events.

**Fig. 6.**
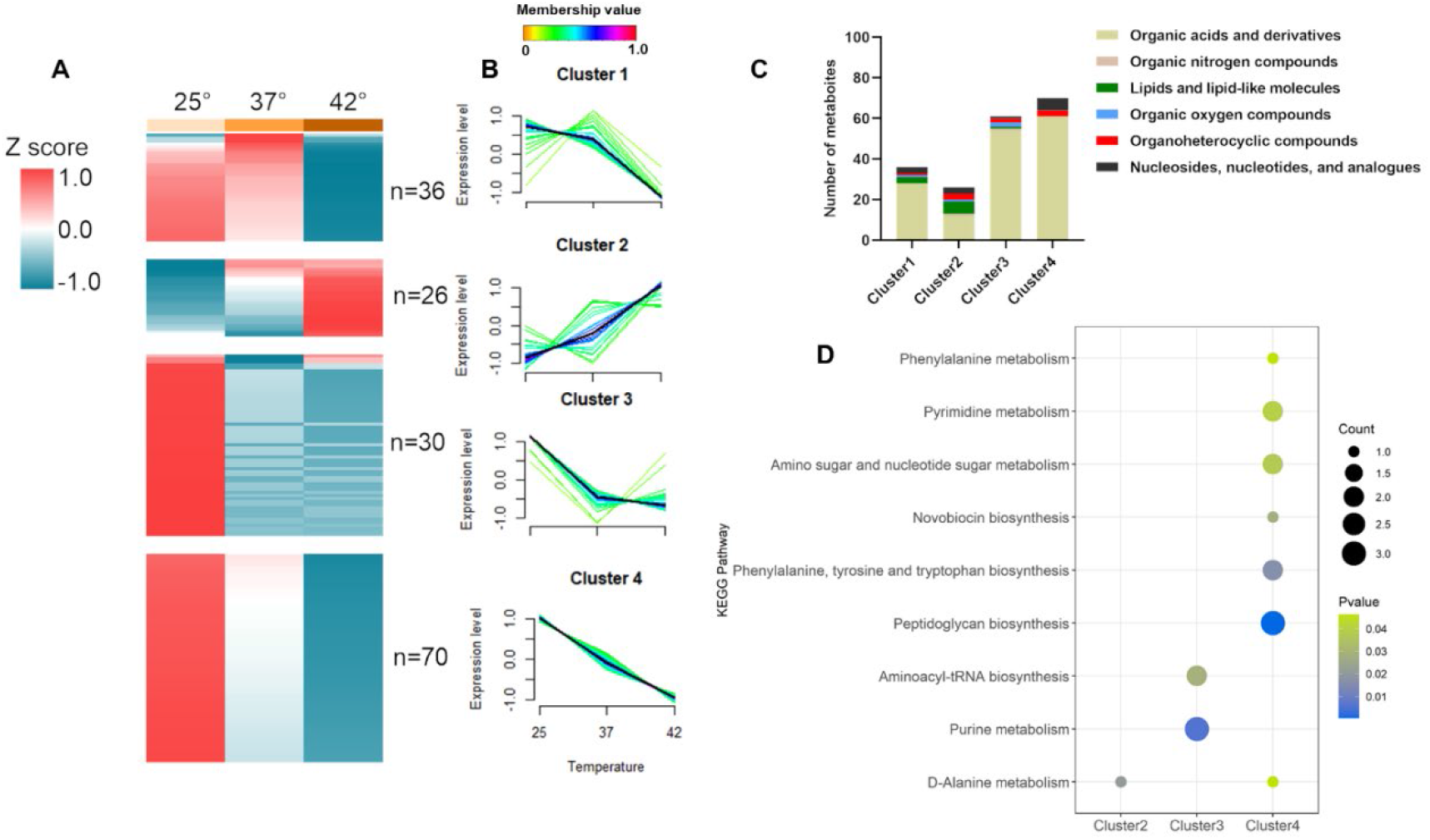
Dynamics of differentially changed metabolites induced by sporulation temperature. Heat map shows the measured relative level of metabolites in spores formed at 42, 37 and 25℃ (A). Mfuzz c-means clustering grouped expression profiles of metabolites (B) Chemical category classification of metabolites in each cluster (C). Bubble chart of fractions of cluster enriched to KEGG pathways (D).

#### Sporulation temperature shapes the unique composition of the *B. subtilis* spore multilayered coat

As the spore coat has also been implicated in temperature resistance (Abhyankar et al., 2016; Kanaan et al., 2022), we next set out to perform an integrated analysis of the proteomic and metabolomic data for the *B. subtilis* spores produced at the three different temperatures (25°C, 37°C, and 42°C) to gain insight into the global response of this spore specific multilayered structure to varying temperature conditions during sporulation. Our results demonstrate that exposure to high temperatures (42°C) led to a distinct composition of protein and metabolites compared to the spores produced at lower temperatures (25°C and 37°C). The thicker spore outer layer cortex is comprised of PG which is similar to that of vegetative cell wall PG (Tipper and Linnett, 1976). Analysis of the spore multilayered coat revealed significant differences in the levels of proteins and metabolites involved in PG biosynthesis, such as UDP-GlcNAc, UDP-MurNAc, and D-Ala-D-Ala, which were found to be decreased in the 42°C spores (Fig. 7). However, D-Ala showed an increase in abundance. We also observed significant increases in the levels of MurF, UppP, and Ddl in the 42°C spores, while MurAB and SpoVD proteins were only higher in the 25°C spores. Furthermore, elevated morphogenetic coat proteins (SpoVID and SpoIVA) as well as CotJC, OxdD and Tgl level were observed in 42°C compared to the low temperatures, while the level of SpoVM and YjzB was decreased in spore basement layer. Additionally, YxeE, YjqC and YybL decreased in the heat stress spores, whereas exhibiting an increase in GerQ and YaaH. In the outer coat, 8 proteins including CotA, CotB, CotC, CotG, CotO, CotQ, CotR and SpsB showed decreased levels at 42°C. CotW is the only crust protein we detected significantly increased in 42°C spores. Other coat proteins, such as Ldt, Ypep, YabG, Yqft, YheC, YvdO and CotSA also showed significant changes but the localization of these have yet to be determined.The presence of CotA, CotB, CotG, and CotQ in the proteomics samples is consistent with previous data (Isticato et al., 2020; Melly et al., 2002).

**Fig. 7.**
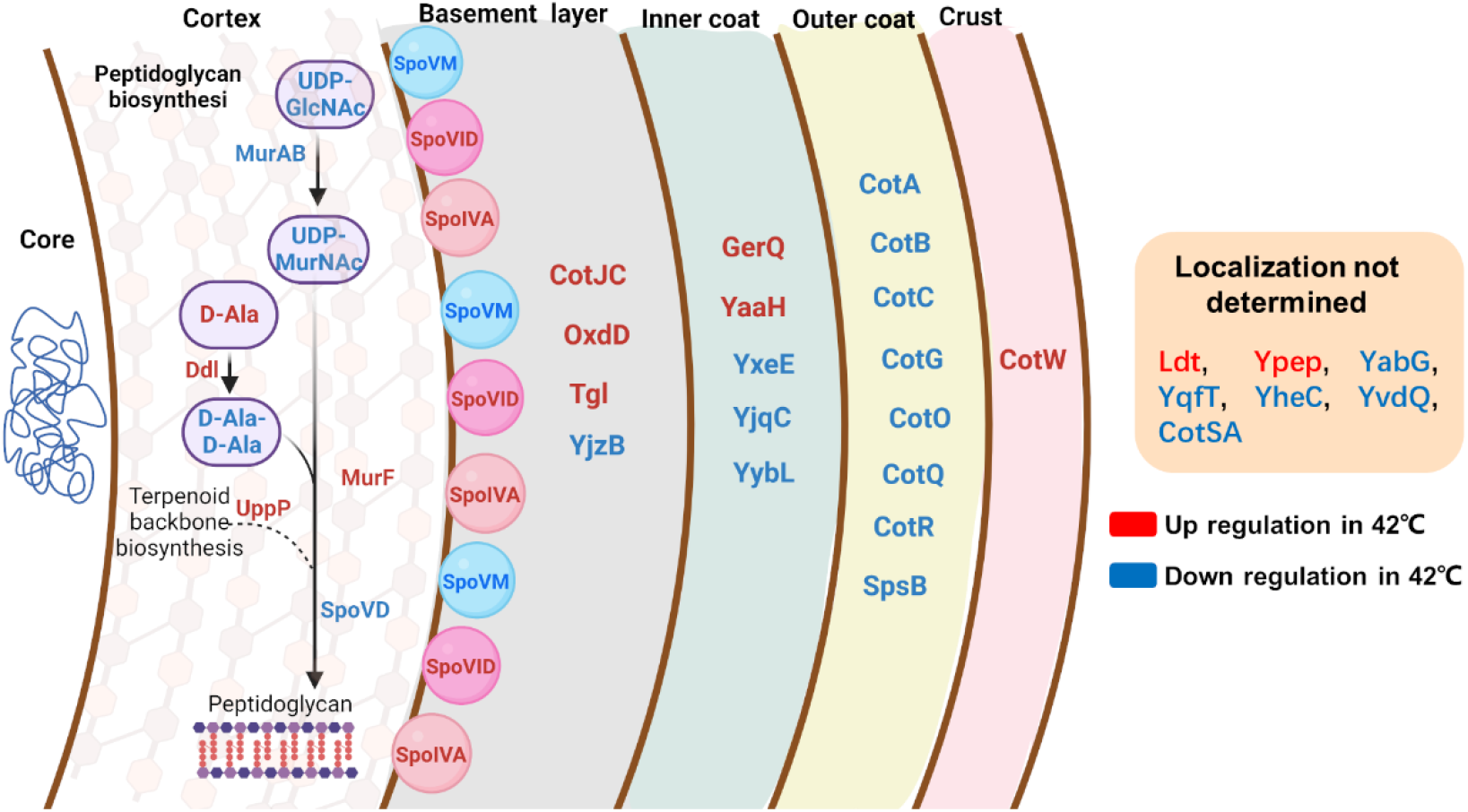
Schematic diagram of coat protein and PG biosynthesis of changes in *B. subtilis* spore multi- layers. Red and blue letters indicate upregulation and downregulation, respectively, in spores grown at 42℃ compared to 25 or 37℃.

## Discussion

The ability to endure extreme growth environments is a fundamental disparity between vegetative cells and spores. In addition to facilitating the survival of vegetative cells, spores also exhibit distinct structural and characteristic features that are shaped by diverse growth conditions. Spores exhibit remarkable sensitivity to environmental changes, either physical or chemical, and adapt accordingly. Among the various environmental factors, the temperature during sporulation constitutes a pivotal determinant of bacterial spore characteristics (Bressuire-Isoard et al., 2018). Consistent with previous studies (Isticato et al., 2020; Melly et al., 2002), our investigation reveals that spores produced under higher environmental temperature (42°) exhibit increased heat resistance and DPA content, as well as a delayed response to AGFK and L-valine as a germination agent. As early as the late 1920s, it was observed that *Bacillus anthracis* spores formed at 37°C exhibited greater heat resistance than those formed at 18°C (Williams, 1929), highlighting the early observation of the impact of environmental conditions on sporulation. More recent research has revealed that the spore’s resistance to wet heat can be attributed to various factors such as the DNA-saturated by α/β-type SASPs, low core water, high DPA and mineral content, as well as alterations in the spore’s structure, which preserve molecular mobility in the core and prevent protein denaturation and irreversible aggregation caused by thermal stress (Abhyankar et al., 2015; Sanchez-Salas et al., 2011; Setlow, 2016; Sunde et al., 2009). Our investigation has revealed that three SASPs, SspJ, SspE, and SspA, along with four HSPs DnaK, GroES, HtpG, and GroEL, were upregulated at 42°C, suggesting that these proteins might have an involvement in acquiring higher heat resistance. Despite previous evidence suggesting that heat shock proteins do not influence spores’ wet heat resistance (Melly and Setlow, 2001), the upregulation of heat shock proteins in response to elevated temperatures has been shown in several studies (Hecker and Völker, 1998; Völker et al., 1994).

At the transcriptional level, sporulation is governed by four sporulation-specific sigma factors, σF, σE, σG, and σK, which are activated post-translationally in specific compartments at specific times (Meeske et al., 2016; Wang et al., 2006). During the early stage of sporulation, σF and σE respectively regulate over 400 important genes in the forespore and mother cell to ensure proper development of key spore elements (Ramos-Silva et al., 2019). Inactivation of the structural genes of σF or σE results in an “abortively disporic” phenotype (Errington, 1993). During the middle stage of sporulation, σG takes over from σF to regulate spore maturation, thus, the rapid generation of spores at higher temperatures could be attributed to the expedited assembly of spores by accelerating the synthesis of crucial components. σK, the last activated sigma factor, plays a critical role in driving the final stages of spore protective structure assembly (cortex and coat) in the mother cell. Notably, we observed a significant reduction in proteins regulated by σK in spores produced at 42°C compared to those generated at 25°C or 37°C. Previous studies have shown that spore coat structure produced at 42°C exhibits a granular, rather than compact and thick outer coat, similar to that of the cotG null mutant strain (Isticato et al., 2020). In addition, we also found that the shape of the spores produced at 42° is also different from the spores produced at 25°, showing a narrower rod-like structure. These observations suggest that these defective structures in spores generated at 42°C may be predominantly due to perturbed regulation of the σK mediated spore developmental phase.

It has been reported that *B. subtilis* modulates its sporulation efficiency to achieve a trade-off between spore quality and quantity, thereby impacting its fitness in diverse environments (Mutlu et al., 2020). In the present study, while the sporulation efficiency was high at 42°C, coat-deficient spores containing 17 proteins with reduced abundance were produced. It also suggests that the suboptimal quantitative levels of spores generated at 25°C may be mitigated by superior quality, which in turn facilitates spore revival (Fig. 1C). Remarkably our proteomics results showed increased levels of most germination proteins in 42°C spores compared to those of 25°C. This observation can be understood in light of prior research indicating that the upregulation of functional germinant receptors (GRs) encoded by GerB and GerK did not result in enhanced germination efficacy with AGFK (Cabrera-Martinez et al., 2003). Given that germination is somehow associated with spore coat proteins, prior data have demonstrated that the absence or mutation of CotG or CotH can impede the L-alanine and AGKF pathways of nutritional germination (Freitas et al., 2020; Isticato et al., 2020; Nguyen et al., 2016). Accordingly, it is possible that spores with analogous coat defects under 42° C have an adverse effect on nutrient uptake. Additionally, enhanced cross-linking of coat proteins might potentially prolong the germination time (Abhyankar et al., 2015). Also noteworthy is our discovery of a significant increase in Tgl within 42° C spores, a protein known to be involved in cross-linking of a few coat proteins (Fernandes et al., 2019; Henriques et al., 1997; Ragkousi and Setlow, 2004; Ursem et al., 2021; Zilhão et al., 2004).

Some rod-shaped bacteria undergo a change in morphology during the process of sporulation. This is particularly evident in Firmicutes such as *Bacilli* and *Clostridia*, where the morphological transformation is from a rod-shaped vegetative cell to an oval-shaped spore (Zhang et al., 2020). Here, we found that spores generated at outside soil environmental temperature of 25°C displayed a spherical morphology in contrast to the rod-shaped spores observed at 37°C and 42°C. Our investigation further revealed that the spores generated at 42°C exhibited a higher abundance of shape-related proteins such as RodZ, YubA, and Mbl, while the levels of RpeA were reduced when compared to spores produced at 25°C. Research has revealed that the morphogenetic protein RodZ and the actin paralogue protein Mbl can interact directly to maintain the rod-shaped morphology of *B. subtilis* (Muchová et al., 2013). In addition, downregulation of RodZ expression can directly lead to the development of cells with a shorter and rounder shape, and mutations in the Mbl gene can also impact cell width in *B. subtilis* (Jones et al., 2001; Muchová et al., 2013). We clearly understand the ultimate distinction between spore and cell, but our data implicitly point towards a possible effect of cell morphology proteins on spore shape. While our knowledge of spore shape formation is limited, it is observed that the absence of Ssdc results in a rounder spore shape (Luhur et al., 2020). This morphological alteration is primarily initiated by its attachment to the spore coat, subsequently impacting the organization of the spore cortex. The PG cell wall is a defining feature of cell geometry in most rod-shaped bacteria, where factors involved in the synthesis and degradation of its major structures are critical in determining cell shape (van Teeffelen and Renner, 2018). The metabolomics results suggested there were 70 spore metabolites present at lower levels in spores formed at stress temperatures versus those formed at 25 °C. These metabolites were significantly enriched with those with a prime role in PG biosynthesis. Specifically, the levels of UDP-GlcNAc, UDP-MurNAc, and D-Ala-D-Ala and enzymes such as MurAB and SpoVD, which are involved in the synthesis of PG were significantly reduced in spores subjected to formation at 42°C. Nevertheless, prior investigations have revealed an increase in cortex muramic acid, that enhances peptidoglycan chain cross-linking as the sporulation temperature increases from 22 to 48°C (Melly et al., 2002). These findings suggest a possible compensatory mechanism for maintaining spore cortex stability in spores, whereby the PG biosynthesis pathway is downregulated while cross-linking is increased.

The coat of *B. subtilis* is comprised of a minimum of 80 distinct proteins. This proteinaceous layer serves a multitude of functions, including but not limited to protection, regulation of germination, and interactions with the external environment. Fig. 7 shows that the level of initiating coat assembly morphogenetic coat proteins SpoIVA, SpoVID and SpoVM were influenced by sporulation temperatures. Except for SpoIVA and SpoVID, CotJC, OxdD and Tgl were more abundant in the basement layer in 42°C spore. GerQ, undergoes cross-linking through the enzymatic action of transglutaminase Tgl and research has demonstrated that Tgl functions in conjunction with YabG to facilitate the temperature-responsive modification of coat protein (Kuwana et al., 2006). Specifically, the cross-linking of GerQ at 37°C is contingent upon the presence of YabG. Importantly, the level of YabG was decreased in spores formed at 42°C. Another important germination protein YaaH that contributes to peptidoglycan hydrolysis is more abundant in 42°C spores. It is noteworthy that a significant reduction in the levels of 8 outer coat proteins, including CotA, CotB, CotC, CotG, CotO, CotQ, CotR, and SpsB, was observed in spores generated during exposure to 42°C. In the crust layer, only CotW was found significantly increased in 42°C spores. The reduced expression of outer layer coat proteins at 42°C may be due to changes in the metabolic state of sporulation or the induction of misfolding or aggregation of coat proteins caused by the increased temperature, leading to their degradation or decreased stability.

## Conclusion

This study provides further integrative metabolic and protein-based evidence that the sporulation temperature significantly affects spore characteristics, such as wet heat resistance, germination response, morphology changes, protein expression, and metabolite composition. Notably, the observed significant differences in PG synthesis and the abundance of spore coat protein components require additional biochemical functional investigation. Overall, this study provides insights into the mechanistic effects that a key environmental factor such as the temperature at spore formation has on the resulting spore properties and highlights the complex adaptation mechanisms governing sporulation in *B. subtilis*. The data can have important implications for the development of strategies to control bacterial spores in various settings, including food preservation, healthcare, and biodefence.

## Acknowledgments

We thank prof. Ezio Ricca for kindly providing the mutation strains used in this study.

## Author contributions

Conceptualization, Y.H., SB and GK; methodology, Y.H.; formal analysis, Y.H.; writing—original draft preparation, Y.H.; writing—review and editing, S.B., and G.K.; resources, W.R. All authors have read and agreed to the published version of the manuscript.

## Data Availability

All mass spectral data have been deposited into Massive (https://massive.ucsd.edu/ProteoSAFe/static/massive.jsp), with the identifier MSV000092146

## Supplementary Data

Table S1 Time of appearance of the fluorescence signal at the various temperatures.

Table S2 Summary of quantitated proteins across all the samples.

Table S3 Differentially expressed proteins among different groups.

Table S4 Summary of annotated metabolites across all the samples.

Table S5 Differentially expressed metabolites among different groups.

## Supplementary figures

**Fig. S1.**
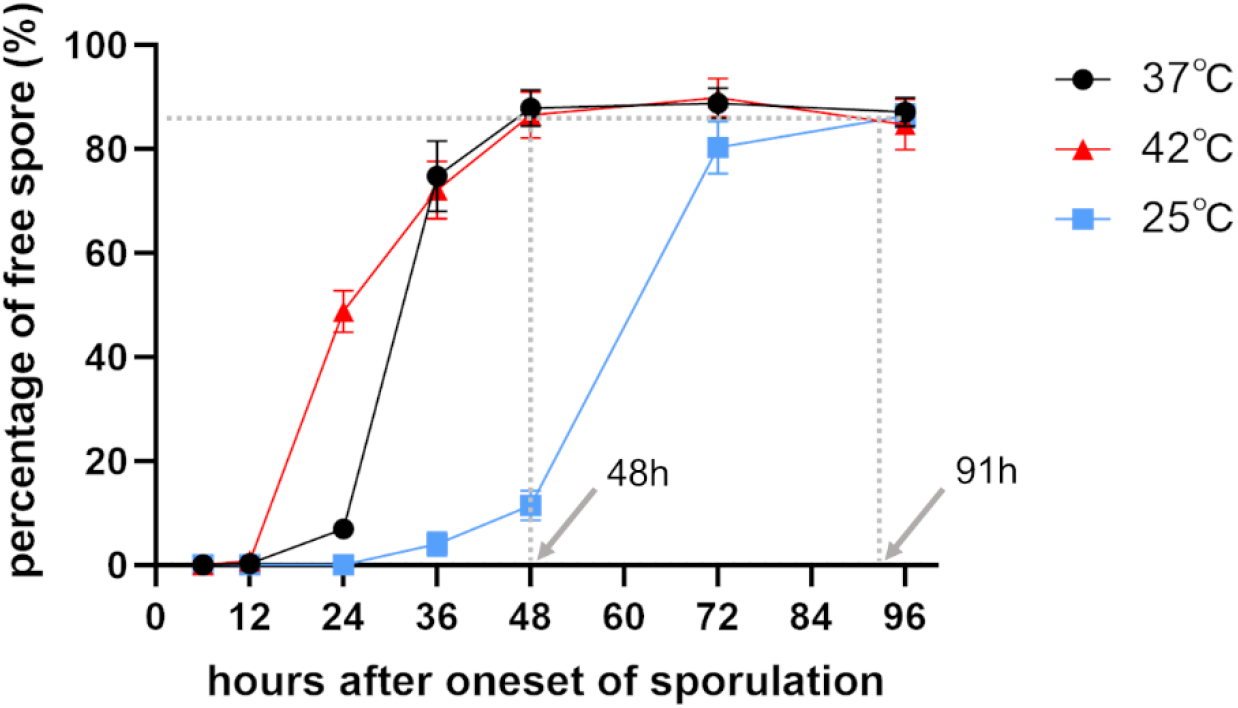
Percentage of free spores during sporulation at 42, 37 and 25℃.

**Fig. S2.**
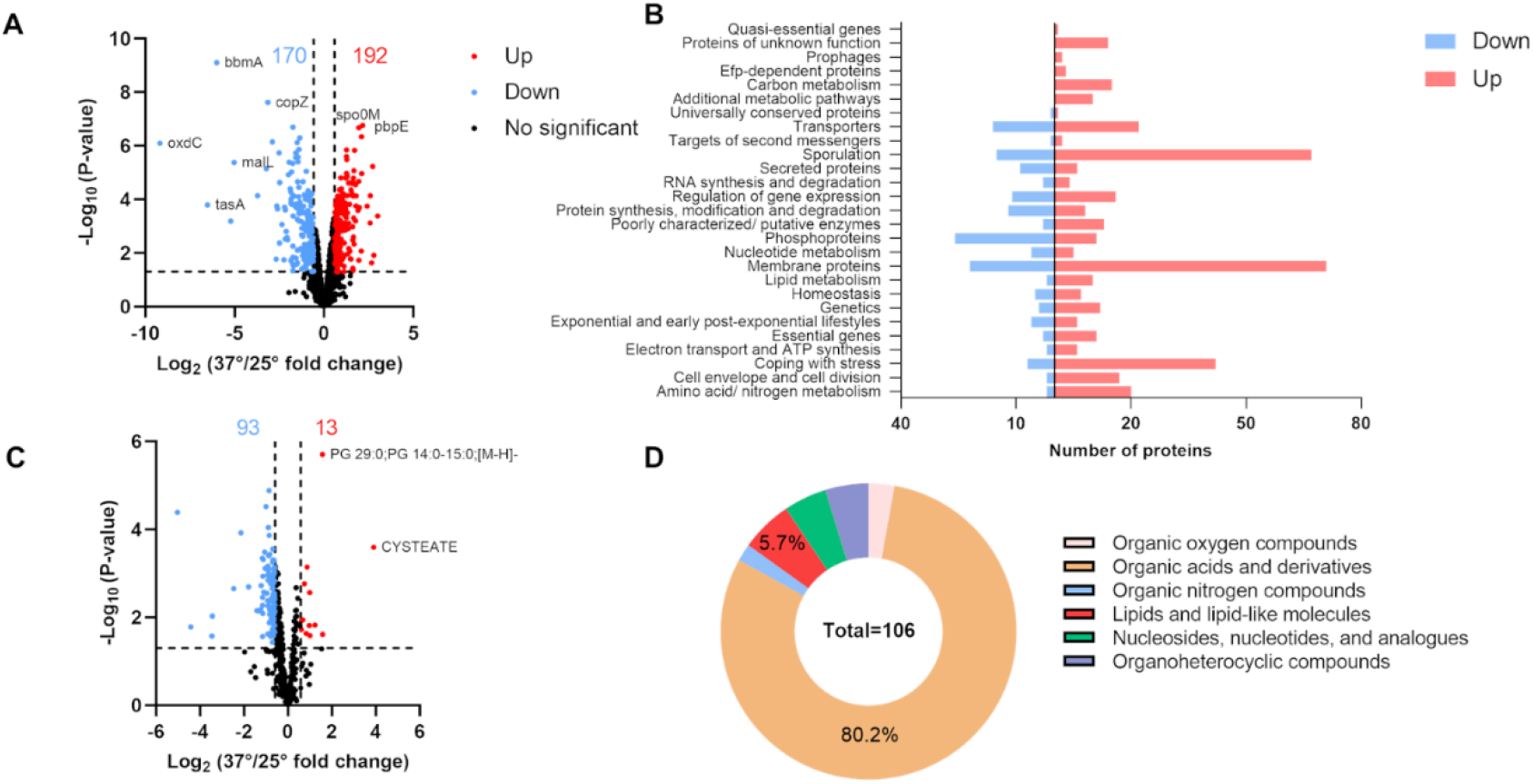
Volcano plots of differential proteins (A) and metabolites (C) in 37 vs. 25℃. Category classification of differentially expressed proteins according to SubtiWiki (B). Chemical category classification of differential metabolites (D).

**Fig. S3.**
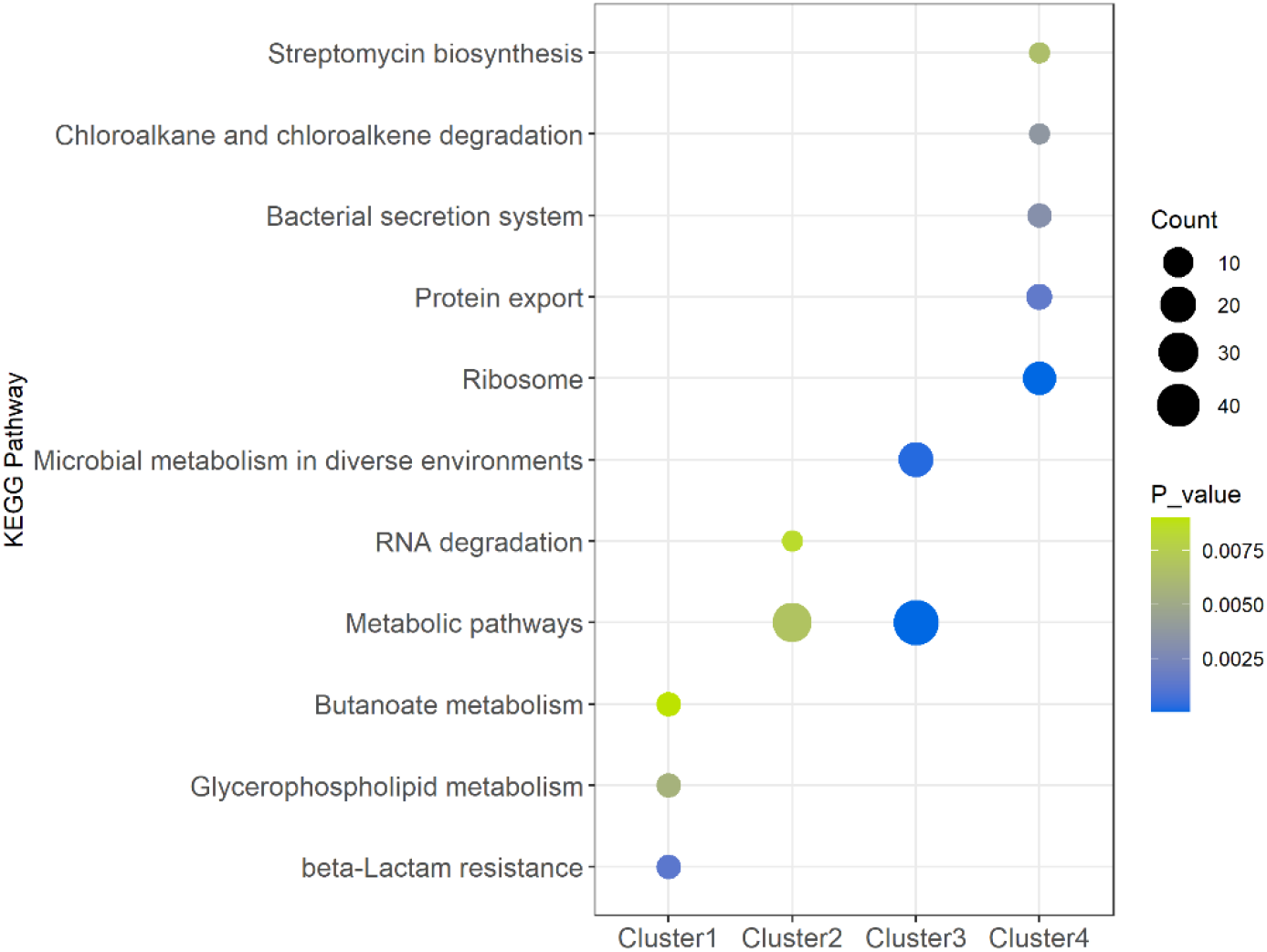
Bubble chart of fractions of cluster enriched to KEGG pathway.

**Fig. S4.**
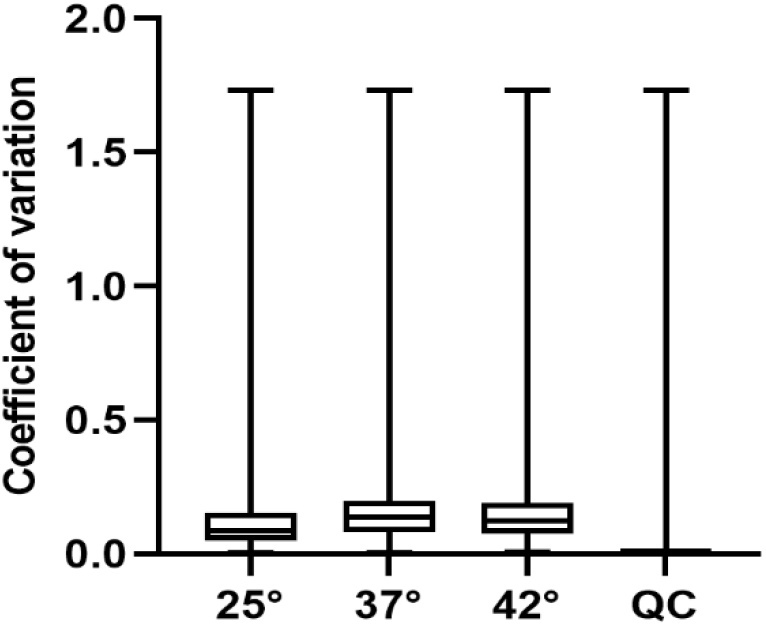
Box plots of CV of the metabolites quantified in at least 2 of 3 replicates in each temperature and QC sample.

## References

Abhyankar, W., Pandey, R., Ter Beek, A., Brul, S., de Koning, L.J., de Koster, C.G., 2015. Reinforcement of Bacillus subtilis spores by cross-linking of outer coat proteins during maturation. Food Microbiol., Spoilers, wonder spores and diehard microorganisms: New insights to integrate these super foes in food spoilage risk management 45, 54–62. https://doi.org/10.1016/j.fm.2014.03.007

Abhyankar, W.R., Kamphorst, K., Swarge, B.N., van Veen, H., van der Wel, N.N., Brul, S., de Koster, C.G., de Koning, L.J., 2016. The Influence of Sporulation Conditions on the Spore Coat Protein Composition of Bacillus subtilis Spores. Front. Microbiol. 7.

Baril, E., Coroller, L., Couvert, O., El Jabri, M., Leguerinel, I., Postollec, F., Boulais, C., Carlin, F., Mafart, P., 2012. Sporulation boundaries and spore formation kinetics of Bacillus spp. as a function of temperature, pH and aw. Food Microbiol. 32, 79–86. https://doi.org/10.1016/j.fm.2012.04.011

Bressuire-Isoard, C., Broussolle, V., Carlin, F., 2018. Sporulation environment influences spore properties in Bacillus: evidence and insights on underlying molecular and physiological mechanisms. FEMS Microbiol. Rev. 42, 614–626. https://doi.org/10.1093/femsre/fuy021

Cabrera-Martinez, R.-M., Tovar-Rojo, F., Vepachedu, V.R., Setlow, P., 2003. Effects of Overexpression of Nutrient Receptors on Germination of Spores of Bacillus subtilis. J. Bacteriol. 185, 2457–2464. https://doi.org/10.1128/JB.185.8.2457-2464.2003

Carobene, A., Braga, F., Roraas, T., Sandberg, S., Bartlett, W.A., 2013. A systematic review of data on biological variation for alanine aminotransferase, aspartate aminotransferase and γ-glutamyl transferase. Clin. Chem. Lab. Med. CCLM 51, 1997–2007. https://doi.org/10.1515/cclm-2013-0096

Checinska, A., Paszczynski, A., Burbank, M., 2015. Bacillus and Other Spore-Forming Genera: Variations in Responses and Mechanisms for Survival. Annu. Rev. Food Sci. Technol. 6, 351–369. https://doi.org/10.1146/annurev-food-030713-092332

Cox, J., Mann, M., 2008. MaxQuant enables high peptide identification rates, individualized p.p.b.-range mass accuracies and proteome-wide protein quantification. Nat. Biotechnol. 26, 1367–1372. https://doi.org/10.1038/nbt.1511

Donadio, G., Lanzilli, M., Sirec, T., Ricca, E., Isticato, R., 2016. Localization of a red fluorescence protein adsorbed on wild type and mutant spores of Bacillus subtilis. Microb. Cell Factories 15, 153. https://doi.org/10.1186/s12934-016-0551-2

Donnelly, M.L., Fimlaid, K.A., Shen, A., 2016. Characterization of Clostridium difficile Spores Lacking Either SpoVAC or Dipicolinic Acid Synthetase. J. Bacteriol. 198, 1694–1707. https://doi.org/10.1128/JB.00986-15

Edwards-Hicks, J., Mitterer, M., Pearce, E.L., Buescher, J.M., 2020. Metabolic Dynamics of In Vitro CD8+ T Cell Activation. Metabolites 11, 12. https://doi.org/10.3390/metabo11010012

Errington, J., 1993. Bacillus subtilis sporulation: regulation of gene expression and control of morphogenesis. Microbiol. Rev. 57, 1–33.

Fernandes, C.G., Martins, D., Hernandez, G., Sousa, A.L., Freitas, C., Tranfield, E.M., Cordeiro, T.N., Serrano, M., Moran, Charles. P., Henriques, A.O., 2019. Temporal and spatial regulation of protein cross-linking by the pre-assembled substrates of a Bacillus subtilis spore coat transglutaminase. PLoS Genet. 15, e1007912. https://doi.org/10.1371/journal.pgen.1007912

Fimlaid, K.A., Shen, A., 2015. Diverse Mechanisms Regulate Sporulation Sigma Factor Activity in the Firmicutes. Curr. Opin. Microbiol. 24, 88–95. https://doi.org/10.1016/j.mib.2015.01.006

Freitas, C., Plannic, J., Isticato, R., Pelosi, A., Zilhão, R., Serrano, M., Baccigalupi, L., Ricca, E., Elsholz, A.K.W., Losick, R., O. Henriques, A., 2020. A protein phosphorylation module patterns the Bacillus subtilis spore outer coat. Mol. Microbiol. 114, 934–951. https://doi.org/10.1111/mmi.14562

Gauvry, E., Mathot, A.-G., Couvert, O., Leguérinel, I., Coroller, L., 2021. Effects of temperature, pH and water activity on the growth and the sporulation abilities of Bacillus subtilis BSB1. Int. J. Food Microbiol. 337, 108915. https://doi.org/10.1016/j.ijfoodmicro.2020.108915

Ghosh, S., Korza, G., Maciejewski, M., Setlow, P., 2015. Analysis of Metabolism in Dormant Spores of Bacillus Species by 31P Nuclear Magnetic Resonance Analysis of Low-Molecular-Weight Compounds. J. Bacteriol. 197, 992–1001. https://doi.org/10.1128/JB.02520-14

Hecker, M., Völker, U., 1998. Non-specific, general and multiple stress resistance of growth-restricted Bacillus subtilis cells by the expression of the sigmaB regulon. Mol. Microbiol. 29, 1129–1136. https://doi.org/10.1046/j.1365-2958.1998.00977.x

Henriques, A.O., Beall, B.W., Moran, C.P., 1997. CotM of Bacillus subtilis, a member of the alpha-crystallin family of stress proteins, is induced during development and participates in spore outer coat formation. J. Bacteriol. 179, 1887–1897.

Hilbert, D.W., Piggot, P.J., 2004. Compartmentalization of Gene Expression during Bacillus subtilis Spore Formation. Microbiol. Mol. Biol. Rev. 68, 234–262. https://doi.org/10.1128/MMBR.68.2.234-262.2004

Hughes, C.S., Foehr, S., Garfield, D.A., Furlong, E.E., Steinmetz, L.M., Krijgsveld, J., 2014. Ultrasensitive proteome analysis using paramagnetic bead technology. Mol. Syst. Biol. 10, 757. https://doi.org/10.15252/msb.20145625

Isticato, R., Lanzilli, M., Petrillo, C., Donadio, G., Baccigalupi, L., Ricca, E., 2020. Bacillus subtilis builds structurally and functionally different spores in response to the temperature of growth. Environ. Microbiol. 22, 170–182. https://doi.org/10.1111/1462-2920.14835

Isticato, R., Scotto Di Mase, D., Mauriello, E.M.F., De Felice, M., Ricca, E., 2007. Amino terminal fusion of heterologous proteins to CotC increases display efficiencies in the *Bacillus subtilis* spore system. BioTechniques 42, 151–156. https://doi.org/10.2144/000112329

Jones, L.J.F., Carballido-López, R., Errington, J., 2001. Control of Cell Shape in Bacteria: Helical, Actin-like Filaments in Bacillus subtilis. Cell 104, 913–922. https://doi.org/10.1016/S0092-8674(01)00287-2

Kanaan, J., Murray, J., Higgins, R., Nana, M., DeMarco, A.M., Korza, G., Setlow, P., 2022. Resistance properties and the role of the inner membrane and coat of Bacillus subtilis spores with extreme wet heat resistance. J. Appl. Microbiol. 132, 2157–2166. https://doi.org/10.1111/jam.15345

Kort, R., O’Brien, A.C., van Stokkum, I.H.M., Oomes, S.J.C.M., Crielaard, W., Hellingwerf, K.J., Brul, S., 2005. Assessment of Heat Resistance of Bacterial Spores from Food Product Isolates by Fluorescence Monitoring of Dipicolinic Acid Release. Appl. Environ. Microbiol. 71, 3556–3564. https://doi.org/10.1128/AEM.71.7.3556-3564.2005

Kumar, L., E. Futschik, M., 2007. Mfuzz: A software package for soft clustering of microarray data. Bioinformation 2, 5–7.

Kuwana, R., Okuda, N., Takamatsu, H., Watabe, K., 2006. Modification of GerQ Reveals a Functional Relationship between Tgl and YabG in the Coat of Bacillus subtilis Spores. J. Biochem. (Tokyo) 139, 887–901. https://doi.org/10.1093/jb/mvj096

Liu, G., Nie, R., Liu, Y., Mehmood, A., 2022. Combined antimicrobial effect of bacteriocins with other hurdles of physicochemic and microbiome to prolong shelf life of food: A review. Sci. Total Environ. 825, 154058. https://doi.org/10.1016/j.scitotenv.2022.154058

Luhur, J., Chan, H., Kachappilly, B., Mohamed, A., Morlot, C., Awad, M., Lyras, D., Taib, N., Gribaldo, S., Rudner, D.Z., Rodrigues, C.D.A., 2020. A dynamic, ring-forming MucB / RseB-like protein influences spore shape in Bacillus subtilis. PLOS Genet. 16, e1009246. https://doi.org/10.1371/journal.pgen.1009246

Meeske, A.J., Rodrigues, C.D.A., Brady, J., Lim, H.C., Bernhardt, T.G., Rudner, D.Z., 2016. High-Throughput Genetic Screens Identify a Large and Diverse Collection of New Sporulation Genes in Bacillus subtilis. PLOS Biol. 14, e1002341. https://doi.org/10.1371/journal.pbio.1002341

Melly, E., Genest, P.C., Gilmore, M.E., Little, S., Popham, D.L., Driks, A., Setlow, P., 2002. Analysis of the properties of spores of Bacillus subtilis prepared at different temperatures. J. Appl. Microbiol. 92, 1105–1115. https://doi.org/10.1046/j.1365-2672.2002.01644.x

Melly, E., Setlow, P., 2001. Heat Shock Proteins Do Not Influence Wet Heat Resistance of Bacillus subtilis Spores. J. Bacteriol. 183, 779–784. https://doi.org/10.1128/JB.183.2.779-784.2001

Miller, R.A., Kent, D.J., Watterson, M.J., Boor, K.J., Martin, N.H., Wiedmann, M., 2015. Spore populations among bulk tank raw milk and dairy powders are significantly different. J. Dairy Sci. 98, 8492–8504. https://doi.org/10.3168/jds.2015-9943

Mou, Y., Mukte, S., Chai, E., Dein, J., Li, X.-J., 2020. Analyzing Mitochondrial Transport and Morphology in Human Induced Pluripotent Stem Cell-Derived Neurons in Hereditary Spastic Paraplegia. J. Vis. Exp. JoVE. https://doi.org/10.3791/60548

Mtimet, N., Trunet, C., Mathot, A.-G., Venaille, L., Leguérinel, I., Coroller, L., Couvert, O., 2015. Modeling the behavior of Geobacillus stearothermophilus ATCC 12980 throughout its life cycle as vegetative cells or spores using growth boundaries. Food Microbiol. 48, 153–162. https://doi.org/10.1016/j.fm.2014.10.013

Muchová, K., Chromiková, Z., Barák, I., 2013. Control of Bacillus subtilis cell shape by RodZ. Environ. Microbiol. 15, 3259–3271. https://doi.org/10.1111/1462-2920.12200

Mutlu, A., Kaspar, C., Becker, N., Bischofs, I.B., 2020. A spore quality–quantity tradeoff favors diverse sporulation strategies in Bacillus subtilis. ISME J. 14, 2703–2714. https://doi.org/10.1038/s41396-020-0721-4

Nguyen, K.B., Sreelatha, A., Durrant, E.S., Lopez-Garrido, J., Muszewska, A., Dudkiewicz, M., Grynberg, M., Yee, S., Pogliano, K., Tomchick, D.R., Pawłowski, K., Dixon, J.E., Tagliabracci, V.S., 2016. Phosphorylation of spore coat proteins by a family of atypical protein kinases. Proc. Natl. Acad. Sci. U. S. A. 113, E3482–E3491. https://doi.org/10.1073/pnas.1605917113

Omardien, S., Ter Beek, A., Vischer, N., Montijn, R., Schuren, F., Brul, S., 2018. Evaluating novel synthetic compounds active against Bacillus subtilis and Bacillus cereus spores using Live imaging with SporeTrackerX. Sci. Rep. 8, 9128. https://doi.org/10.1038/s41598-018-27529-4

Pandey, R., Ter Beek, A., Vischer, N.O.E., Smelt, J.P.P.M., Brul, S., Manders, E.M.M., 2013. Live cell imaging of germination and outgrowth of individual bacillus subtilis spores; the effect of heat stress quantitatively analyzed with SporeTracker. PloS One 8, e58972. https://doi.org/10.1371/journal.pone.0058972

Planchon, S., Dargaignaratz, C., Levy, C., Ginies, C., Broussolle, V., Carlin, F., 2011. Spores of Bacillus cereus strain KBAB4 produced at 10°C and 30°C display variations in their properties. Food Microbiol. 28, 291–297. https://doi.org/10.1016/j.fm.2010.07.015

Postollec, F., Mathot, A.-G., Bernard, M., Divanac’h, M.-L., Pavan, S., Sohier, D., 2012. Tracking spore-forming bacteria in food: From natural biodiversity to selection by processes. Int. J. Food Microbiol. 158, 1–8. https://doi.org/10.1016/j.ijfoodmicro.2012.03.004

Ragkousi, K., Setlow, P., 2004. Transglutaminase-Mediated Cross-Linking of GerQ in the Coats of Bacillus subtilis Spores. J. Bacteriol. 186, 5567–5575. https://doi.org/10.1128/JB.186.17.5567-5575.2004

Ramos-Silva, P., Serrano, M., Henriques, A.O., 2019. From Root to Tips: Sporulation Evolution and Specialization in Bacillus subtilis and the Intestinal Pathogen Clostridioides difficile. Mol. Biol. Evol. 36, 2714–2736. https://doi.org/10.1093/molbev/msz175

Ritchie, M.E., Phipson, B., Wu, D., Hu, Y., Law, C.W., Shi, W., Smyth, G.K., 2015. limma powers differential expression analyses for RNA-sequencing and microarray studies. Nucleic Acids Res. 43, e47. https://doi.org/10.1093/nar/gkv007

Sanchez-Salas, J.-L., Setlow, B., Zhang, P., Li, Y., Setlow, P., 2011. Maturation of Released Spores Is Necessary for Acquisition of Full Spore Heat Resistance during Bacillus subtilis Sporulation. Appl. Environ. Microbiol. 77, 6746–6754. https://doi.org/10.1128/AEM.05031-11

Setlow, P., 2016. Spore Resistance Properties, in: Driks, A., Eichenberger, P. (Eds.), The Bacterial Spore. ASM Press, Washington, DC, USA, pp. 201–215. https://doi.org/10.1128/9781555819323.ch10

Setlow, P., 2014. Spore Resistance Properties. Microbiol. Spectr. 2, 2.5.11. https://doi.org/10.1128/microbiolspec.TBS-0003-2012

Sunde, E.P., Setlow, P., Hederstedt, L., Halle, B., 2009. The physical state of water in bacterial spores. Proc. Natl. Acad. Sci. 106, 19334–19339. https://doi.org/10.1073/pnas.0908712106

Tipper, D.J., Linnett, P.E., 1976. Distribution of peptidoglycan synthetase activities between sporangia and forespores in sporulating cells of Bacillus sphaericus. J. Bacteriol. 126, 213–221. https://doi.org/10.1128/jb.126.1.213-221.1976

Trunet, C., Mtimet, N., Mathot, A.-G., Postollec, F., Leguerinel, I., Sohier, D., Couvert, O., Carlin, F., Coroller, L., 2015. Modeling the Recovery of Heat-Treated Bacillus licheniformis Ad978 and Bacillus weihenstephanensis KBAB4 Spores at Suboptimal Temperature and pH Using Growth Limits. Appl. Environ. Microbiol. 81, 562. https://doi.org/10.1128/AEM.02520-14

Tyanova, S., Temu, T., Sinitcyn, P., Carlson, A., Hein, M.Y., Geiger, T., Mann, M., Cox, J., 2016. The Perseus computational platform for comprehensive analysis of (prote)omics data. Nat. Methods 13, 731–740. https://doi.org/10.1038/nmeth.3901

Ursem, R., Swarge, B., Abhyankar, W.R., Buncherd, H., de Koning, L.J., Setlow, P., Brul, S., Kramer, G., 2021. Identification of Native Cross-Links in Bacillus subtilis Spore Coat Proteins. J. Proteome Res. 20, 1809–1816. https://doi.org/10.1021/acs.jproteome.1c00025

van Teeffelen, S., Renner, L.D., 2018. Recent advances in understanding how rod-like bacteria stably maintain their cell shapes. F1000Research 7, 241. https://doi.org/10.12688/f1000research.12663.1

Völker, U., Engelmann, S., Maul, B., Riethdorf, S., Völker, A., Schmid, R., Mach, H., Hecker, M., 1994. Analysis of the induction of general stress proteins of Bacillus subtilis. Microbiol. Read. Engl. 140 (Pt 4), 741–752. https://doi.org/10.1099/00221287-140-4-741

Wang, C., Morrissey, E.M., Mau, R.L., Hayer, M., Piñeiro, J., Mack, M.C., Marks, J.C., Bell, S.L., Miller, S.N., Schwartz, E., Dijkstra, P., Koch, B.J., Stone, B.W., Purcell, A.M., Blazewicz, S.J., Hofmockel, K.S., Pett-Ridge, J., Hungate, B.A., 2021. The temperature sensitivity of soil: microbial biodiversity, growth, and carbon mineralization. ISME J. 15, 2738–2747. https://doi.org/10.1038/s41396-021-00959-1

Wang, S.T., Setlow, B., Conlon, E.M., Lyon, J.L., Imamura, D., Sato, T., Setlow, P., Losick, R., Eichenberger, P., 2006. The Forespore Line of Gene Expression in Bacillus subtilis. J. Mol. Biol. 358, 16–37. https://doi.org/10.1016/j.jmb.2006.01.059

Williams, O.B., 1929. The Heat Resistance of Bacterial Spores. J. Infect. Dis. 44, 421–465.

Zhang, H., Mulholland, G.A., Seef, S., Zhu, S., Liu, J., Mignot, T., Nan, B., 2020. Establishing rod shape from spherical, peptidoglycan-deficient bacterial spores. Proc. Natl. Acad. Sci. 117, 14444–14452. https://doi.org/10.1073/pnas.2001384117

Zilhão, R., Serrano, M., Isticato, R., Ricca, E., Moran, C.P., Henriques, A.O., 2004. Interactions among CotB, CotG, and CotH during Assembly of the Bacillus subtilis Spore Coat. J. Bacteriol. 186, 1110–1119. https://doi.org/10.1128/JB.186.4.1110-1119.2004

